# White matter hyperintensities and microplastics

**DOI:** 10.1101/2024.11.26.625277

**Authors:** Elaine L. Bearer, Marcus A. Garcia, Natalie Adolphi, Matthew J. Campen

## Abstract

**Synopsis:** White matter hyperintensities are abnormalities that appear in MRI scans of living patients but are not apparent in MRI post-mortem.

**Goal:** Our goal is to understand the cellular/biological basis of white matter hyperintensities (WMH).

**Approach:** We aligned post-mortem MR scans with those collected ante-mortem and performed histopathology and pyrGC/MS for plastics on regions with WMH.

**Result:** PyrGC/MS detected large amounts of plastics and we determined their cellular locations with a novel optical imaging approach in regions with small vessel disease and Abeta plaques.

**Impact:** Microplastics in the brains of people with cognitive impairment may be due to pre-existing vascular injury or contribute to it. Many questions remain: Where do they come from, do they impair function? Can they be diagnosed by MRI ante-mortem?

## Introduction

Microplastics are turning up everywhere, including in normal human brain^1^. Ante-mortem MRI FLAIR images show white matter hyperintensities (WMH) are associated with small vessel disease and occur in Binswanger’s disease encephalopathy (BSE) with motor dysfunction and cognitive decline but are also common in older adults without these diseases. The cellular basis for this abnormality is unknown. Here we report discovery of abundant microplastic fibrils in human forebrain white matter and choroid plexus in regions of WMH in cases of Alzheimer’s (AD) and BSE.

## Methods

We aligned ante-mortem Flair MR images to images acquired post-mortem of the fixed brain (**Fig. 1**). This allowed us to identify regions that display WMH in life and guide gross pathology descriptions and select the abnormal regions for further histopathological investigation. We compared two cases, one with Binswanger’s disease and large subcortical regions of WMH, and another early onset Alzheimer’s disease. We performed standard neurodegenerative pathological post-mortem work-up and examined histopathology slides microscopically. We selected additional blocks from the abnormal subcortical regions for histopathology. Tissue was also harvested for pyrolysis gas chromatography from adjacent regions within the WMH. A new method was used to detect the plastics and type them by PyGC/MS ^2^.

**Figure 1:**
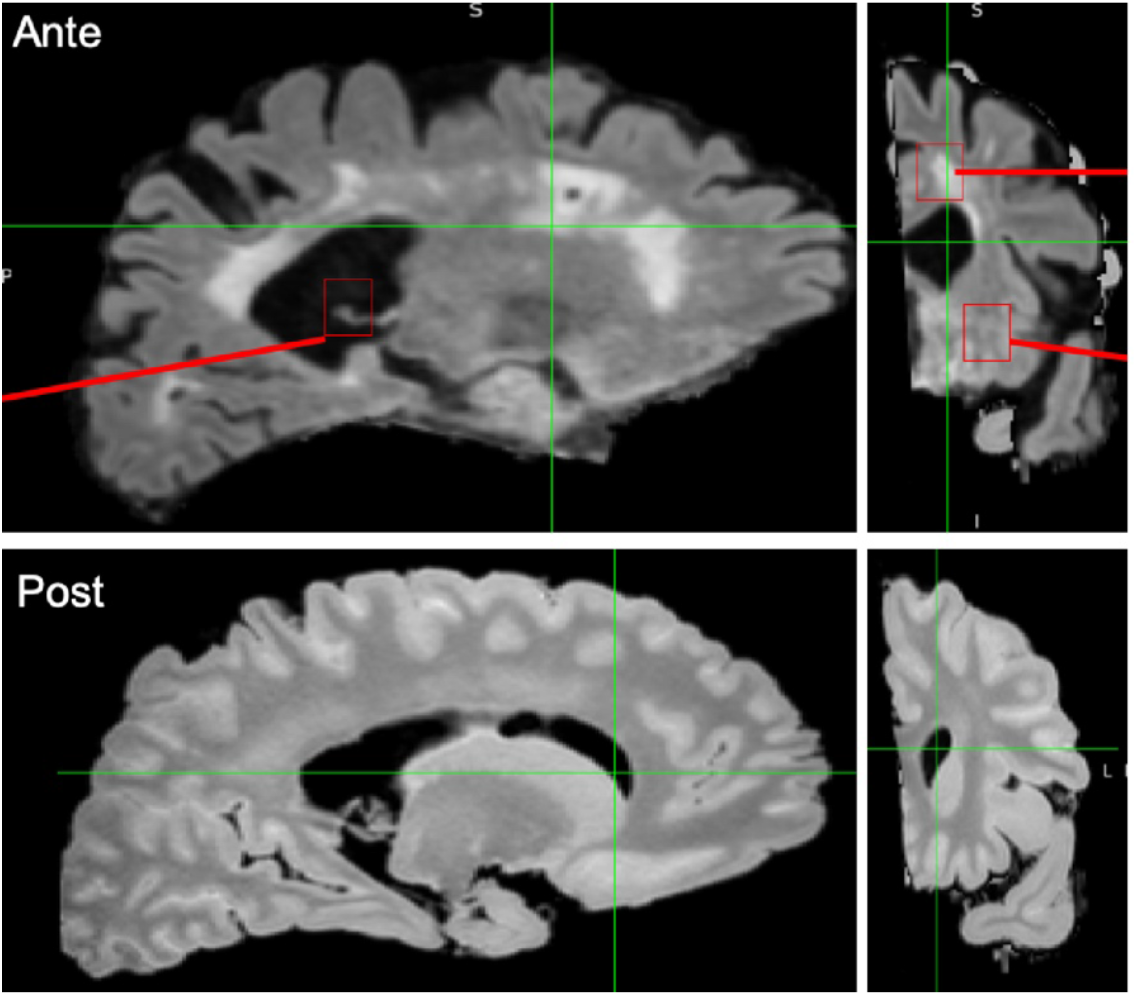
Ante (top panels): FLAIR MR image of a participant in the NM ADRC during clinic visit. Note shrunken gyri and abnormal white subcortical hyperintensities. Red squares indicate location where histopathology samples were taken, in the choroid plexus (left panel), and in the subcortical white matter (right panel, top) and stratum (right panel lower). **Post:** The same brain post-mortem, fixed in formalin and imaged in the same 3T Siemens Sonata MR scanner. Note the hyperintense signal is difficult to appreciate, ventricles are smaller and other anatomy is altered after fixatoin.

We developed a new fluorescence method to detect the microplastics in sections of brain using typical hematoxylin-eosin stained diagnostic slides. First we prepared isolated particles from each brain. Then we scanned the particles in a multi-laser confocal scanning microscope (FV4000, Evident) with 10 different laser lines for excitation and 8 detector wave lengths for emissions. Then we performed a lambda scan for emission spectra for a set of isolated particles, and finally performed Z-stack imaging of individual particles together with a background image using the excitation/emission profile that yielded the best profile separate from background with the isolated particles. Then we imaged H&E stained slides of choroid plexus and cortical regions from post-mortem brains of cases with Binswanger’s or Alzheimer’s diseases. Finally, we examined the isolated microplastics by negative-stain electron-microscopy.

## Results

WMH disappeared in FLAIR images of formalin-fixed post-mortem brain, and gross pathology examination of fixed brain slices noted no evident abnormality. By aligning ante-mortem images with those collected post-mortem, we were able to identify regions that we abnormal in life with the cut brain and select those regions for further study, both chemical and histological (**Fig. 1**).

Microscopy confirmed that the AD case was positive for ABeta plaques, p-tau and neuritic plaques (CERAD) while Binswanger’s had only loose Abeta not in plaques and little to no p-tau. Neither disease had appreciable TDP43 or synuclein aggregates. The BSE case had microhemorrhages in cortical white matter and minimal arteriolar sclerosis. Clusters of CD68 positive macrophages, loss of myelin and rarefication in forebrain white matter corresponded to regions of subcortical WMH by FLAIR MRI, not visible in MR of post-mortem brains. Histopathology showed unusual glossy deposits near blood vessels in regions with myelin loss and abundant macrophages negative for activated microglial markers.

Analysis for microplastics by pyrolysis–gas chromatography/mass spectrometry (PyGC/MS) after alkali treatment and solvent extraction revealed a concentration of 30,908µg/g (ug plastic per g of tissue) in white matter of AD, and 21,441µg/g in BSE–an order of magnitude above that found in normal brains (average, 4,800µg/g)^1^. Cortical gray matter from both diseases also contained more microplastics than normal brains. Fluorescent imaging of isolated microplastics from the brain showed clusters of curling nanofibrils also seen after negative staining in EM (**Fig. 2**). Fluorescent particles with similar characteristics were detected in the choroid plexuses, gray matter and white matter from both AD and BSE. In AD, the particles seemed aggregated within the brain, while in BSE they were more diffuse.

**Figure 2.**
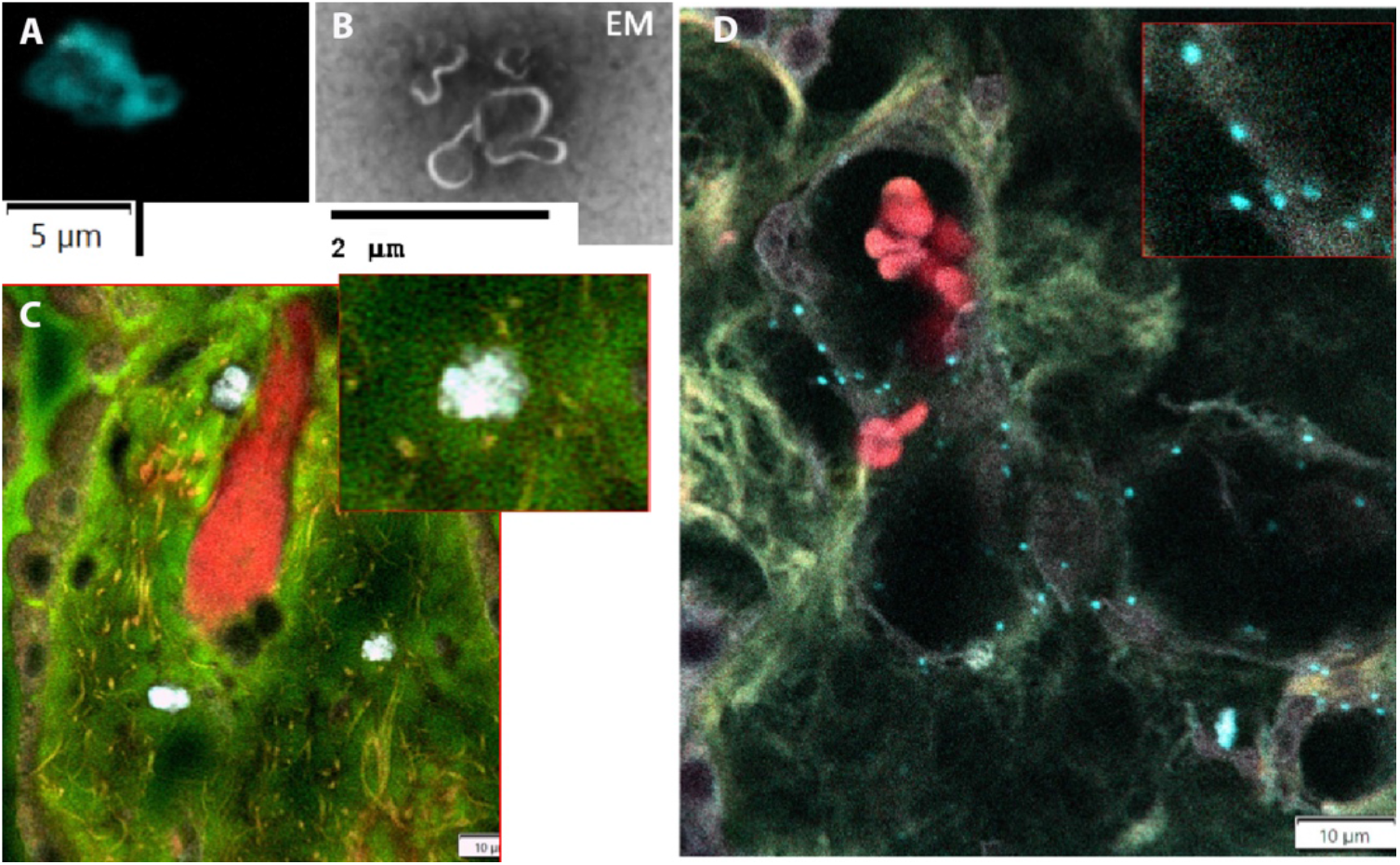
Confocal and EM of microplastics in brain of cognitively impaired adults. **A**. Isolated micro-plastics from the brain of the participant shown in Fig. 1, imaged by confocal microscopy at Caltech. B. Electron-microscopy of these same isolated micro, nano-plastics.. C. Microplastics in the choroid plexus (pale blue) of a case of AD. Connective tissue (green) and red blood cells (red). D. Microplastics in a case of BSE, with the same fluorescence spectra optics as the isolated microplastics.

Negative stain electron-microscopy of isolated plastics revealed curly nano-fibrils, less than 2 nm in diameter, varying in length from 0.5um to 5um or longer from the BSE case. Some were corkscrew-like, most had sharp tips that could pierce blood vessel walls. Many were embedded in aggregates of apparently tissue-derived material, as the brains were fixed in formalin prior to extraction and thus microplastics not well solubilized. We are exploring chemical, optical and ultrastructural analyses with various immuno-histology and heavy metal stains to detect the cellular context of these nanofibers and identify the types of plastics they represent.

## Conclusions

Many questions remain, such as the consequences of such non-biological material in the brain on endothelial cell integrity and blood brain barrier leakage, on activation of inflammatory mediators (matrix metalloproteinases), on macrophages and microglia, as well as on neurons and other glia, and their contribution to impairments of cognitive and motor functions. Can microplastics in brain be diagnosed ante-mortem by MRI?

## Acknowledgements

We thank Gary Rosenberg and John Adair for following these participants in clinic, Arvind Caprihan for ante-mortem-post-mortem alignments, Karen Santa-Cruz for gross brain dissections, Gabriel Dvorak and Brit Vallejo for assistance with brain extractions and dissections, and Cathleen Martinez for histopathology. We are grateful to Andres Collazo, Director of the Imaging Center at the Beckman Institute of California Institute of Technology, Louie Kerr, Director of the Imaging Center at Marine Biology Laboratory in Woods Hole, MA, and Joe DeGiorgis, as well as Alan Gillman, James Lopez and Mark Eastman of Evident/Olympus Scientific. Supported by NIA P30 AG086404 (GR, ELB), NIGMS P20 GM130422 (MJC), and the Harvey Family (ELB). The authors have no competing interests.

